## Supplementary Information for "Analysis of emergent bivalent antibody binding identifies the molecular reach as a critical determinant of SARS-CoV-2 neutralisation potency"

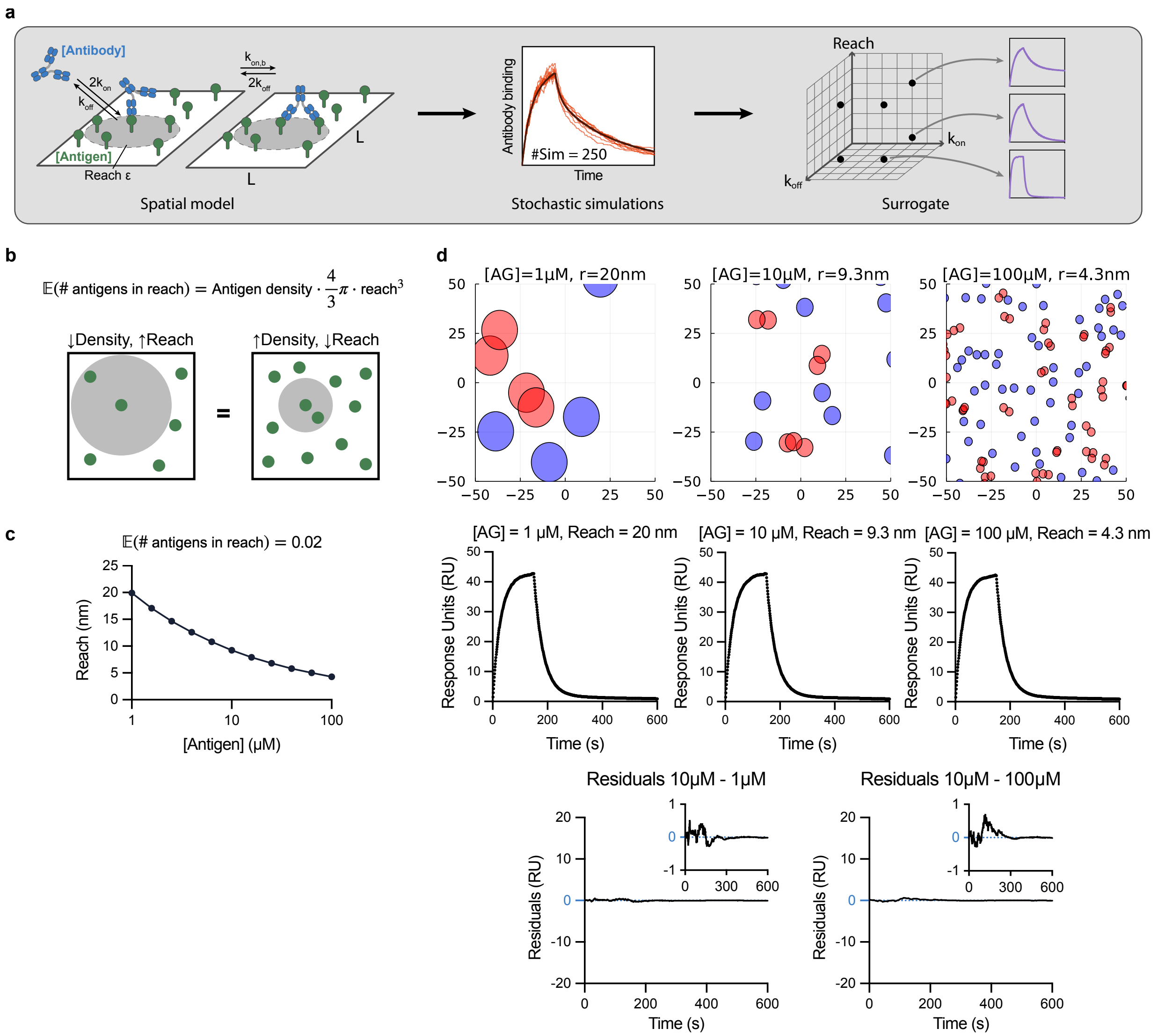


Figure S1: **Workflow for fitting the particle-model to bivalent SPR data.** **(A)** A schematic of the particle-model chemical reactions (left) used to perform simulations of the association and dissociation phase of antibody binding in bivalent SPR (middle, red traces), which are averaged to produce the predicted SPR trace for a given parameter set (middle, black traces). These averaged SPR traces are pre-tabulated for different parameter values ($k_{\text{on}}$, $k_{\text{off}}$, $k_{\text{on,b}}$, reach - $\varepsilon$) using a computing cluster to produce a surrogate for the particle-model that is used in data fitting (right). **(B)** The average number of antigens within reach of an individual antigen depends on the molecular reach and the antigen density. **(C)** The relationship between antigen density and molecular reach when the average number of antigens within reach is 0.02 (equation plotted is from panel b). **(D)** The spatial distribution of antigen (top) and the corresponding predicted bivalent SPR traces (bottom) for different antigen concentrations and molecular reach calculated using the relationship in panel B. The SPR traces are effectively identical as shown by the small residuals (difference between the indicated SPR curves) confirming that antibody binding depends on the average number of antigens within reach, which can be achieved by a short reach at high antigen density or a long reach at low antigen density. Simulations are performed using an antibody concentration of 1 nM with $k_{\text{on}} = 0.05\ \mu$ $\text{M}^{-1}\text{s}^{-1}$, $k_{\text{off}} = 0.02\text{s}^{-1}$, and $k_{\text{on,b}} = 1.0^{-1}$.

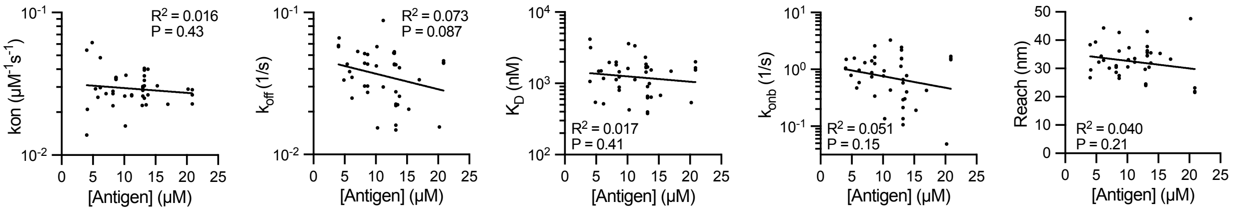


Figure S2: **The fitted binding parameters for the IgG FD-11A antibody binding RBD determined by fitting the particle-model to bivalent SPR data are independent of the RBD antigen density.** The indicated parameter is plotted over the RBD antigen density ($N = 42$). The coefficient of determination ($R^2$) and the p-value for the null hypothesis that the fitted line and a horizontal line (i.e. no relationship between the binding parameter and RBD density) produce an equal fit to the data.

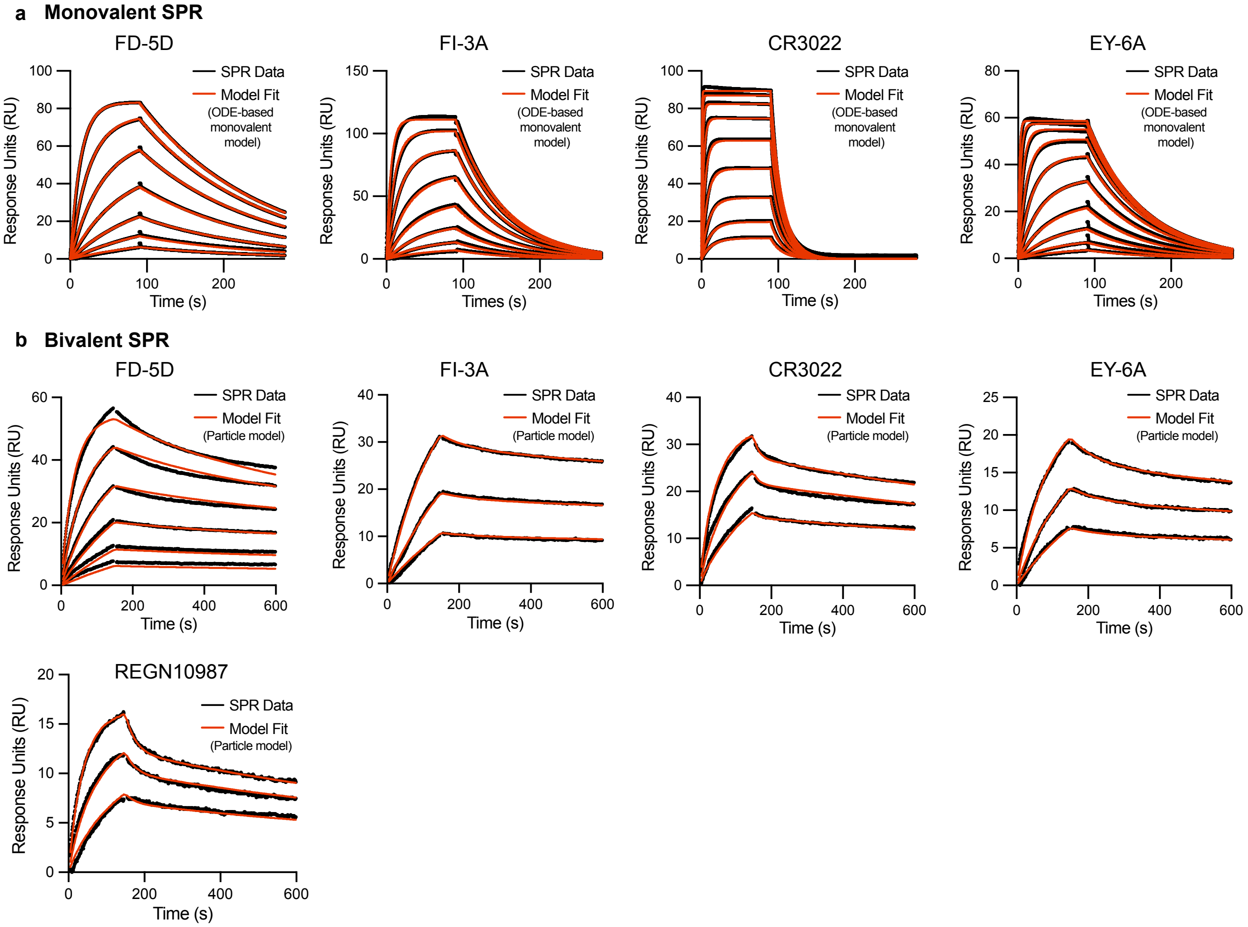


Figure S3: **Representative monovalent and bivalent SPR for the indicated IgG1 antibodies binding RBD. (A)** Representative monovalent SPR traces produced by injecting RBD (2000 nM with 2-fold dilutions) over surfaces immobilised with the indicated antibody. **(B)** Representative bivalent SPR traces produced by injecting the indicated antibodies over surfaces immobilised with RBD. Antibodies were injected using a 2-fold dilution series, with a top concentration of 300 nM, 5 nM, 5 nM, 5 nM, 5 nM for FD-5D, FI-3A, CR3022, EY-6A and REGN10987 respectively. RBD was immobilised at 7 - 15 $\mu$M.

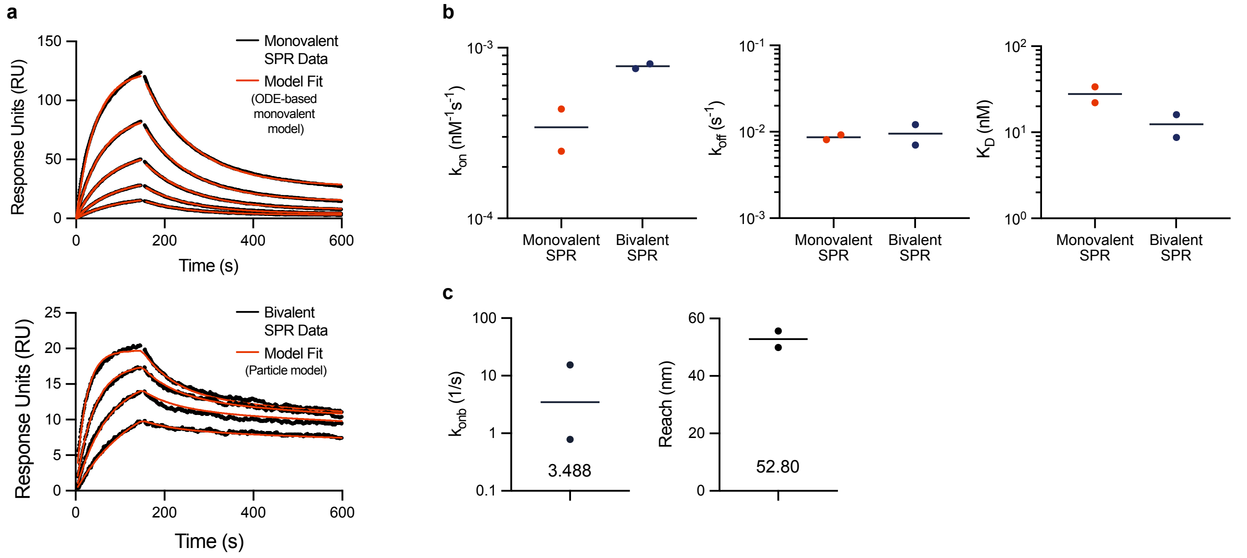


Figure S4: **The particle-model accurately analyses bivalent SPR generated using the CD19 antigen reproducing the monovalent binding parameters.** Data is generated using purified Spytag-CD19 (antigen) and an anti-CD19 antibody (SJ25C1). **(A)** Monovalent SPR (top) is produced by injecting Spytag-CD19 over a surface immobilised with SJ25C1 and bivalent SPR (bottom) is produced by injecting SJ25C1 over a surface immobilised with Spytag-CD19 (through coupling to amine-coupled Spycatcher). The SPR traces are generated by 2-fold dilution of CD19 starting at 175 nM (top) or by a 2-fold dilution of SJ25C1 starting at 33 nM (bottom). Representative SPR traces (black) and model fit (red) are shown for 1 out of 2 representative experiments. **(B)** The indicated parameter produced by monovalent or bivalent SPR. **(C)** The indicated parameter produced by bivalent SPR.

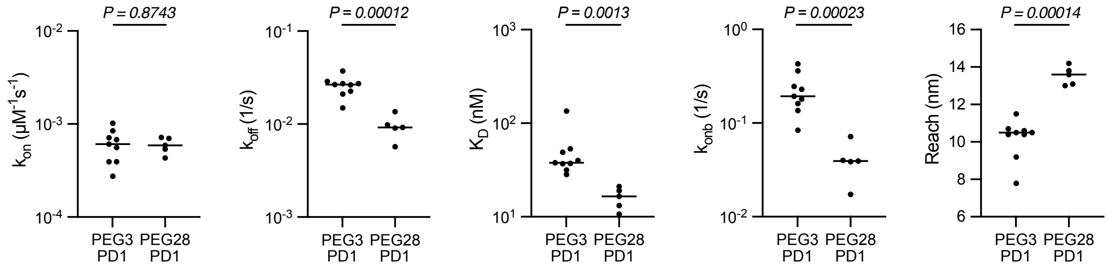


Figure S5: **The fitted binding parameters of an anti-phosphotorysine antibody (PY20) interacting with a small phosphorylated peptide antigen coupled to PEG3 or PEG28 linker.** The indicated parameter from N=9 or N=5 independent experiments for PEG3 and PEG28 coupled phosphorylated peptide antigen, respectively. A t-test with a Holm-Šídák correction for multiple comparison was used to determine p-values.

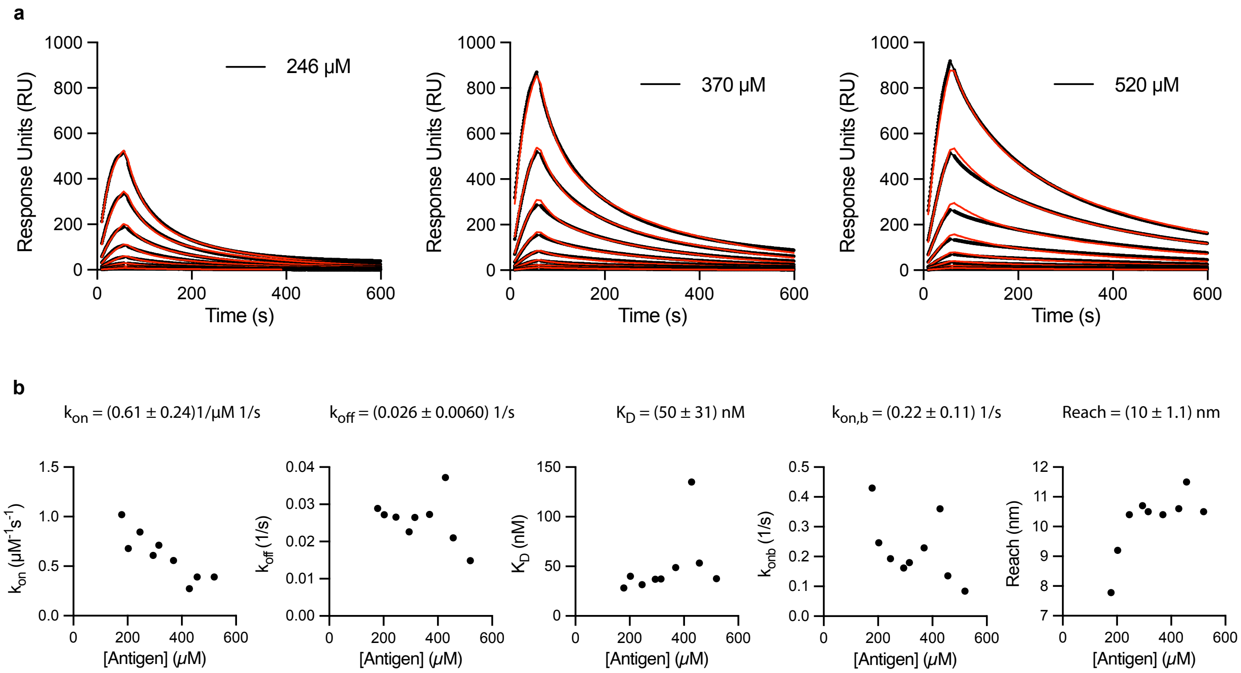


Figure S6: **The fitted binding parameters for an anti-phosphotorysine antibody (PY20) interacting with different densities of a small phosphorylated peptide antigen coupled to PEG3.** **(A)** Representative SPR traces (black) and particle-based model fits (red) for the PY20 antibody injected over surfaces with the indicated concentration of PEG3 coupled to a small phosphorylated peptide antigen. The antibody was injected at 8 different concentrations using a 2-fold dilution from a top concentration of 25 nM. **(B)** The fitted binding parameters plotted over the density of the small phosphorylated peptide antigen coupled to the SPR chip surface.

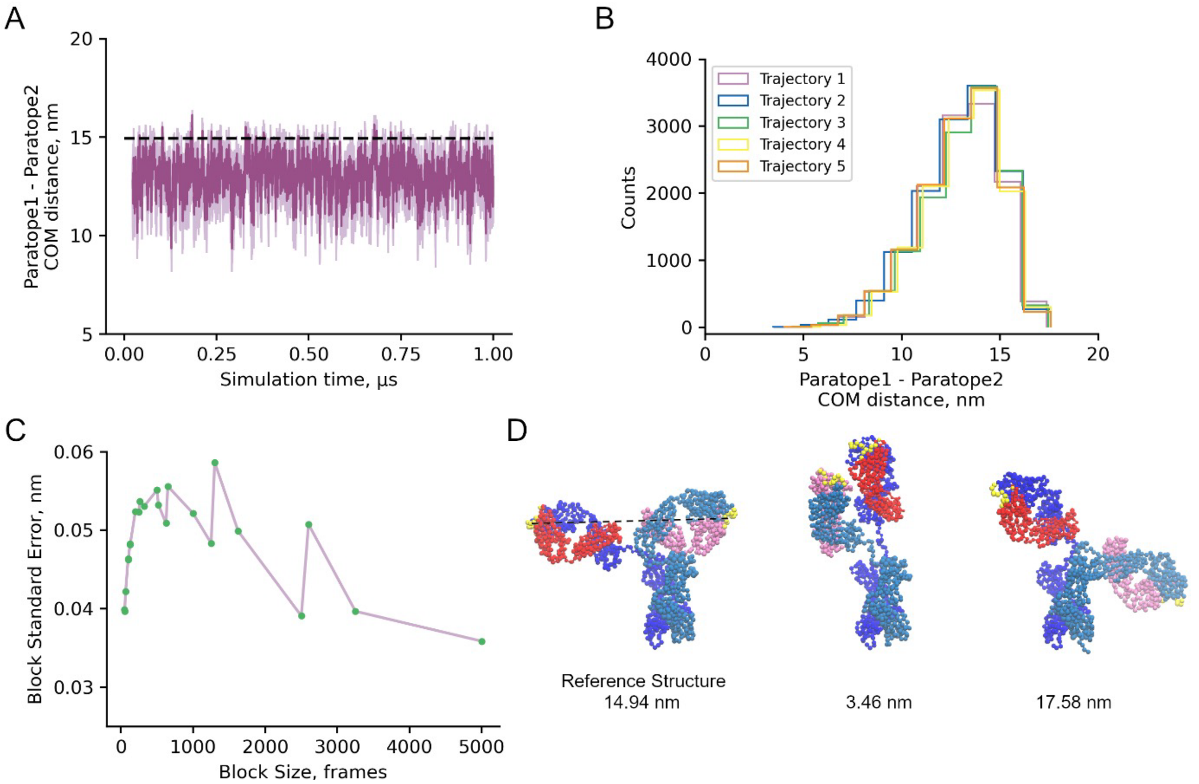


Figure S7: **The mean distance between the antigen-binding domains of the IgG1 FD-11A antibody is 13 nm.** The FD-11A Fab structure is used to generate a coarse-grain model of the complete IgG1 FD-11A antibody (see methods) and is used in coarse-grained MD simulations. **(A)** Ensemble average time series (dark purple) computed over five independent 1-$\mu$s trajectories of the distance between the centres-of-mass of the coarse-grain interaction sites that form the paratopes on each Fab binding arm (i.e. the distance between the antigen binding domains). The light purple shaded region indicates the standard error over the five trajectories at each simulation frame. The dotted black line indicates the distance between the antigen-binding sites in the structure. **(B)** Histogram of the inter-paratope or inter-antigen binding domain distances over each of the five independent trajectories. **(C)** Block averaging analysis of the merged trajectory indicates a block size of 1,000 frames is suitable. The mean inter-paratope distance is 13.05 ± 0.05 nm (error bar is estimated as the block standard error with a block size of 1000 frames). Block averaging was performed on a merged trajectory consisting of the final 975 ns of the five independent runs. **(D)** Coarse-grain structures of the reference structure (left), minimum reach structure from simulations (middle), and maximum reach structure from simulations (right). The two heavy chains are colored dark and light blue, the two light chains are colored red and pink, and the interactions sites constituting the paratopes are colored yellow. The dotted black line in the leftmost structure indicates the distance between the centers-of-mass of the paratopes in the reference state (14.94 nm).

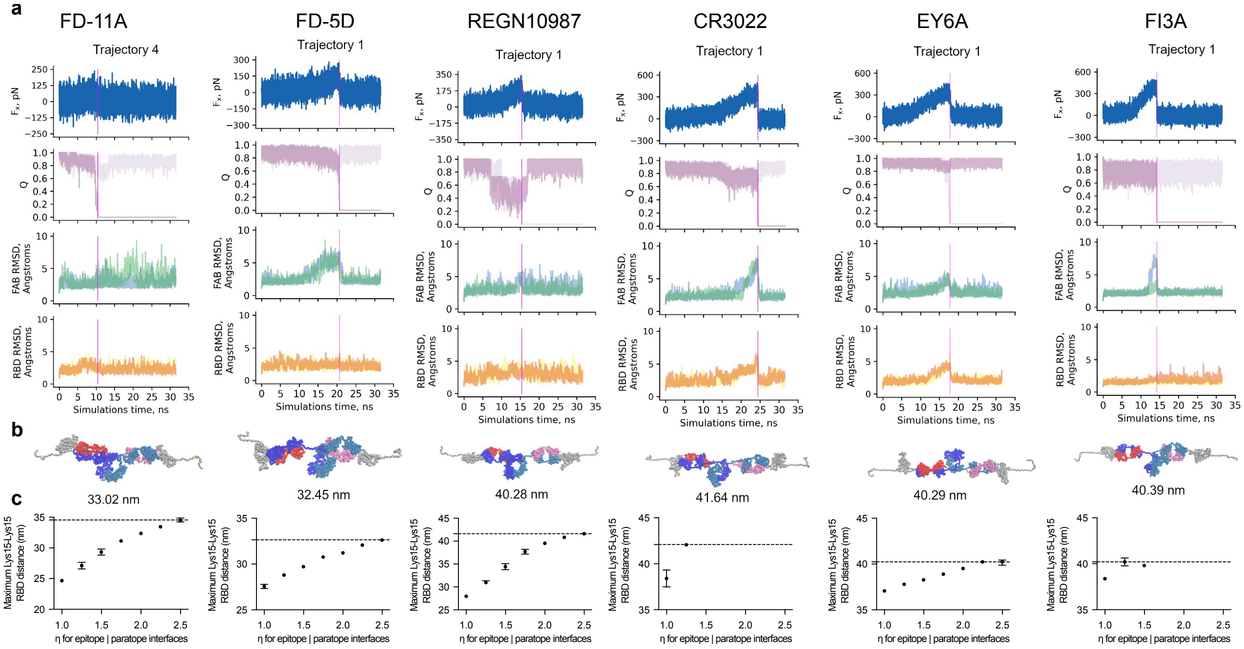


Figure S8: **Estimating the molecular reach of RBD-specific antibodies using coarse-grained molecular dynamic simulations.** **(A)** A representative coarse-grained MD simulation trajectory for the indiciated antibody (columns) showing the calculated force along the pulling direction ($F_x$), the fraction of native contacts for each antigen interface (Q, interface 1 in pink and 2 in mauve), the RMSD as a function of time during the simulation for Fabs 1 and 2 (blue and green, respectively), and the RMSD of RBD 1 and 2 (yellow and orange, respectively). **(B)** The structure from the trajectories in panel A that produced the maximum distance between the Lys15 residues on RBD (shown below the structure) whilst the antibody was bound bivalently. This structure was achieved at the time point indicated by the vertical magenta lines in panel A. **(C)** The maximum Lys15-Lys15 distance over the interface binding strength ($\eta$) calculated over the set of trajectories where unfolding does not take place during the MD simulations. Error bars are 95% confidence intervals computed from bootstrapping with $10^6$ independent samples. Missing data points for CR3022 and FI3A indicate that all trajectories were unfolded. The molecular reach is defined as the maximum Lys15-Lys15 distance over all $\eta$ and indicated by the dashed horizontal line.

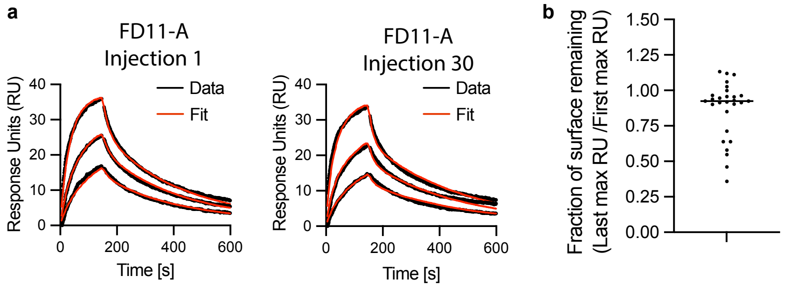


c

| | Passed | Failed | | |
|---|---|---|---|---|
| | Good particle model fit, better than monovalent fit | No regeneration | Poor particle model fit | Good particle model fit, Equal monovalent fit |
| **Count** | 45 | 7 | 12 | 16 |
| **Ab names** | See Fig. 4 | 219, 280, 293, 300, 301, 387 | 2, 334, 379, 412, 270, 340, 264, 253-165, 253-55, 292, 369, 373 | 183, 217, 222, 78, 88, 91, 144, 150, 158, 337, 398, 185, 263, 400 |

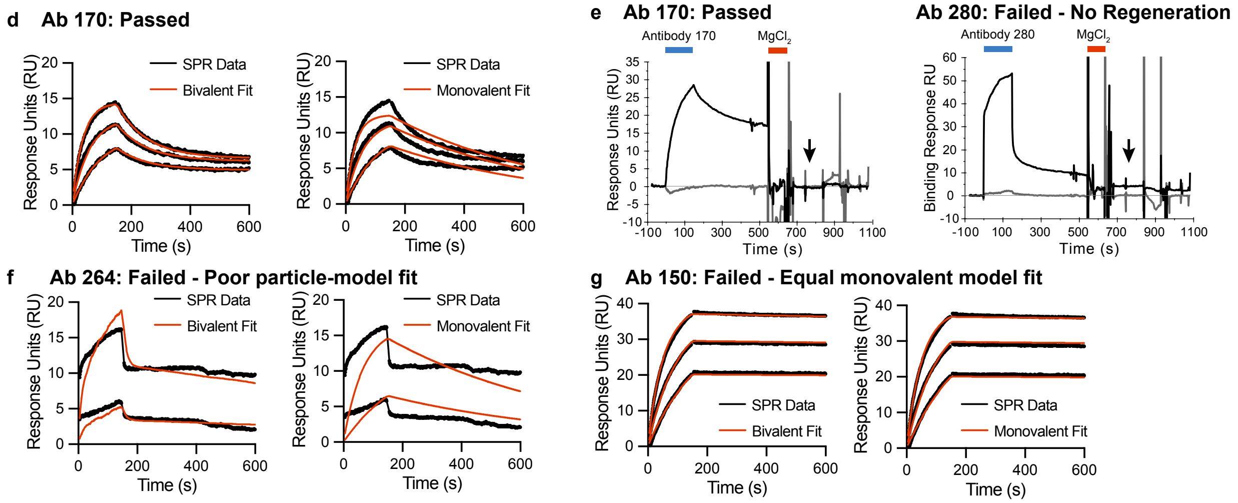


Figure S9: **Quality controls for high throughput analysis of 80 RBD antibodies using bivalent SPR. (A-B)** Surface stability of RBD was assessed by injecting FD-11A at the beginning and end of each experiment. Representative SPR traces of the first and last injection of FD-11A (A) and fraction of FD-11A binding at end of the experiment (B). The fraction is calculated as the RU at 150 s after injection of highest concentration in the first divided by the last injection. **(C)** A summary of the antibodies that passed or failed quality control and hence included or excluded from the analysis, respectively. **(D-G)** Examples of antibodies representing the four possible quality control outcomes. (D) The antibody 170 was included because the particle-model produced a good fit and the monovalent model produced a poor fit. (E) The antibody 170 displays complete regeneration (included) whereas antibody 280 shows only partial regeneration (excluded). Partial regeneration can be observed by residual RU after the injection of 3 M $MgCl_2$ for 90 s at the end of each SPR cycle (see arrow). (F) The antibody 264 was excluded because the particle-model produced a poor fit. (G) The antibody 150 was excluded because the particle-based model (left) and the ODE-based monovalent model (right) produced an equally good fit.

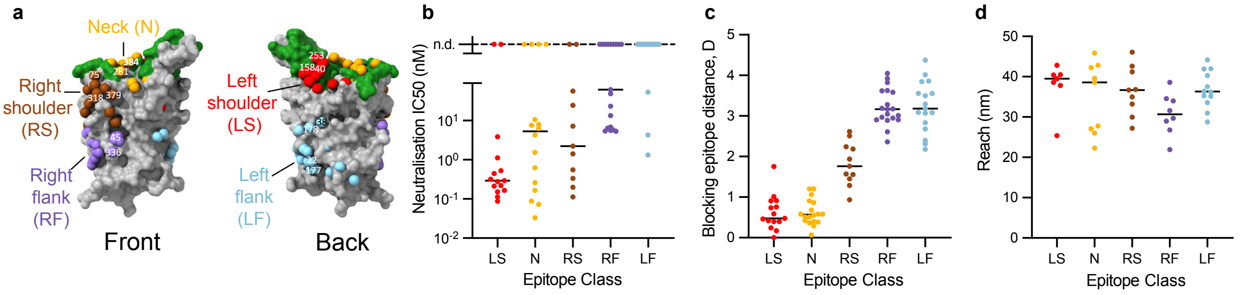


Figure S 10: **The previously introduced epitope taxonomy for RBD-binding antibodies does not stratify the molecular reach.** **(A)** The previously introduced epitope taxonomy for RBD antibodies based on five regions. Figure adapted from (5). **(B-D)** Antibody neutralisation $IC_{50}$ (B), epitope blocking distance (C), and molecular reach of each antibody (D) organised by their taxonomic class.

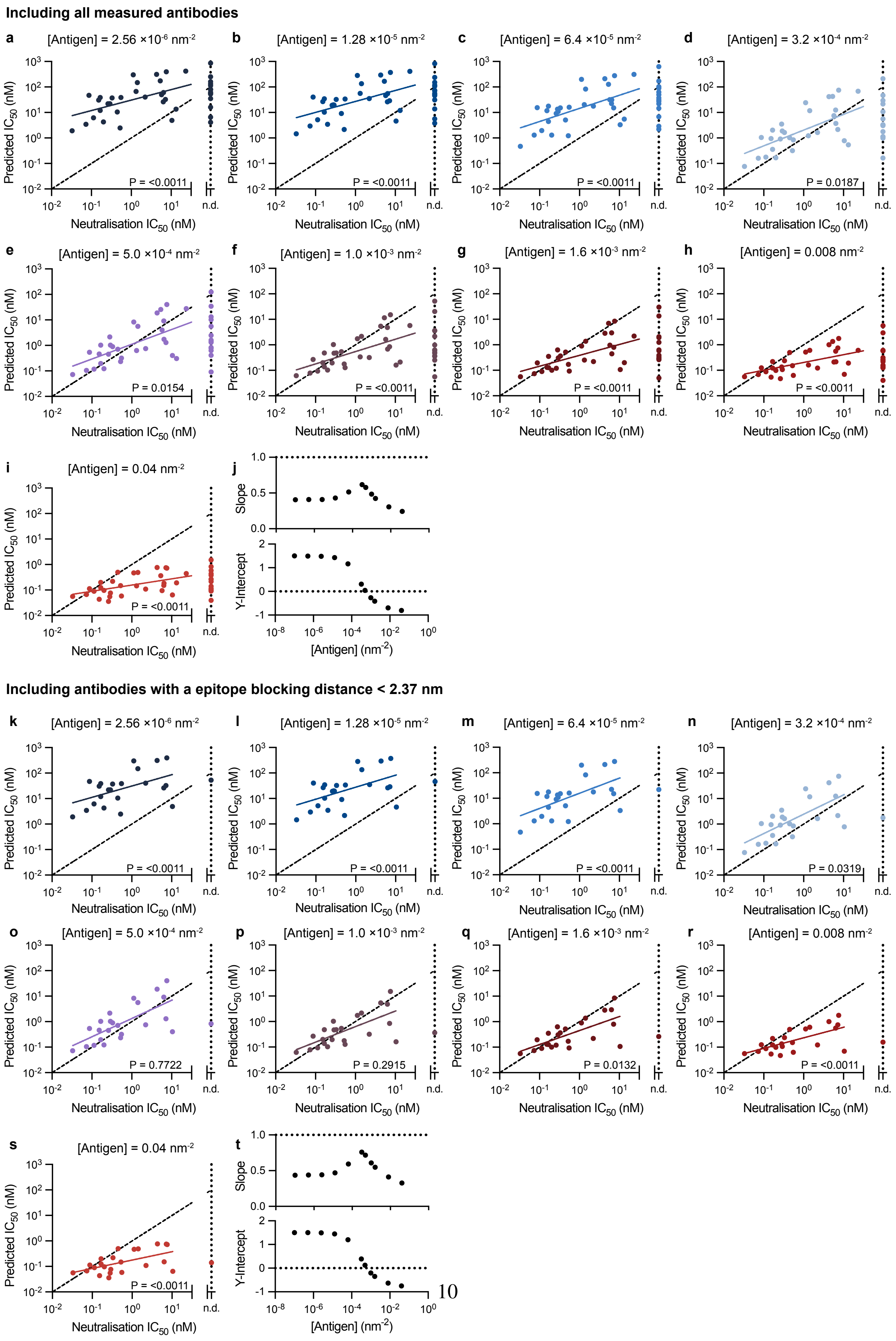
Including all measured antibodies
a [Antigen] = 2.56 ×10-6 nm-2 P = <0.0011
b [Antigen] = 1.28 ×10-5 nm-2 P = <0.0011
c [Antigen] = 6.4 ×10-5 nm-2 P = <0.0011
d [Antigen] = 3.2 ×10-4 nm-2 P = 0.0187
e [Antigen] = 5.0 ×10-4 nm-2 P = 0.0154
f [Antigen] = 1.0 ×10-3 nm-2 P = <0.0011
g [Antigen] = 1.6 ×10-3 nm-2 P = <0.0011
h [Antigen] = 0.008 nm-2 P = <0.0011
i [Antigen] = 0.04 nm-2 P = <0.0011
j
Slope
Y-Intercept
[Antigen] (nm-2)
Predicted IC50 (nM)
Neutralisation IC50 (nM)
n.d.
Including antibodies with a epitope blocking distance < 2.37 nm
k [Antigen] = 2.56 ×10-6 nm-2 P = <0.0011
l [Antigen] = 1.28 ×10-5 nm-2 P = <0.0011
m [Antigen] = 6.4 ×10-5 nm-2 P = <0.0011
n [Antigen] = 3.2 ×10-4 nm-2 P = 0.0319
o [Antigen] = 5.0 ×10-4 nm-2 P = 0.7722
p [Antigen] = 1.0 ×10-3 nm-2 P = 0.2915
q [Antigen] = 1.6 ×10-3 nm-2 P = 0.0132
r [Antigen] = 0.008 nm-2 P = <0.0011
s [Antigen] = 0.04 nm-2 P = <0.0011
t
Slope
Y-Intercept
[Antigen] (nm-2)

Figure S11: **The predicted antibody binding potency produces absolute agreement with the experimental neutralisation potency at intermediate antigen densities.** The particle-based model was used to predict the concentration of antibody required to bind 50% of antigen (Predicted $IC_{50}$) to surfaces randomly distributed with the indicated density of antigen for **(A-I)** all antibodies or **(K-S)** the subset of antibodies that bound within 2.37 nm of the blocking epitope. A linear fit to the log-transformed $IC_{50}$ values (solid line) is compared to a line of absolute agreement (dashed black line - slope 1, y-intercept 0) using an F-test for the null hypothesis that the two lines have the same slope and intercept. A Bonferroni multiple-comparison correction is applied by multiplying each p-value by 11 (number of antigen densities test). **(J,T)** Display the slope (top) and intercept (bottom) of the fitted line for the 11 different antigen densities tested.

Table S1: All-atom model building details

| Antibody | FAB structure source | Chains used (heavy, light, RBD) |
|---|---|---|
| FD-11A | 7PQZ | A, B, E |
| FD-5D | 7PR0 | H, L, E |
| REGN10987 | 6XDG | C, A, E |
| CR3022 | 6W41 | H, L, C |
| EY6A | 6ZDH | H, L, A |
| FI3A | 7PQY | H, L, E |

Table S2: RBD-antibody-RBD chain lengths

| Antibody | Chain | Residues |
|---|---|---|
| FD-5D | HC1 | 1-459 |
| | LC1 | 1-221 |
| | HC2 | 1-459 |
| | LC2 | 1-221 |
| | RBD1 | 1-220 |
| | RBD2 | 1-220 |
| FD-11A | HC1 | 1-457 |
| | LC1 | 1-218 |
| | HC2 | 1-457 |
| | LC2 | 1-218 |
| | RBD1 | 1-220 |
| | RBD2 | 1-220 |
| REGN10987 | HC1 | 1-450 |
| | LC1 | 1-216 |
| | HC2 | 1-450 |
| | LC2 | 1-216 |
| | RBD1 | 1-220 |
| | RBD2 | 1-220 |
| CR3022 | HC1 | 1-449 |
| | LC1 | 1-221 |
| | HC2 | 1-449 |
| | LC2 | 1-221 |
| | RBD1 | 1-220 |
| | RBD2 | 1-220 |
| EY6A | HC1 | 1-451 |
| | LC1 | 1-215 |
| | HC2 | 1-451 |
| | LC2 | 1-215 |
| | RBD1 | 1-220 |
| | RBD2 | 1-220 |
| FI3A | HC1 | 1-447 |
| | LC1 | 1-214 |
| | HC2 | 1-447 |
| | LC2 | 1-214 |
| | RBD1 | 1-220 |
| | RBD2 | 1-220 |

Table S3: $\eta$ values for all antibody and RBD domains and interfaces except the epitope—paratope interfaces. *HC1 and LC2 share a small interface in the 1HZH crystal structure

| Identity | Structural Class | $\eta$ |
|---|---|---|
| HC1 | $\beta$ | 2.480 |
| LC1 | $\beta$ | 2.480 |
| HC2 | $\beta$ | 2.480 |
| LC2 | $\beta$ | 2.480 |
| RBD 1 | $\alpha/\beta$ | 1.916 |
| RBD 2 | $\alpha/\beta$ | 1.916 |
| HC1—LC1 interface | - | 2.124 |
| HC2—LC2 interface | - | 2.124 |
| HC1—HC2 interface | - | 2.124 |
| HC1—LC2 interface* | - | 1.507 |
